# Identification of *Klebsiella pneumoniae* complex members using MALDI-TOF mass spectrometry

**DOI:** 10.1101/350579

**Authors:** Carla Rodrigues, Virginie Passet, Sylvain Brisse

**Author notes:** **Corresponding author:** Sylvain Brisse, Biodiversity & Epidemiology of Bacterial Pathogens, Institut Pasteur, 25 Rue du Docteur Roux, 75015 Paris Cedex 15, France.

## Abstract

*Klebsiella pneumoniae* (phylogroup Kp1), one of the most problematic pathogens associated with antibiotic resistance worldwide, is phylogenetically closely related to *K. quasipneumoniae* [subsp. *quasipneumoniae* (Kp2) and subsp. *similipneumoniae* (Kp4)], *K. variicola* (Kp3) and two unnamed phylogroups (Kp5 and Kp6). Together, Kp1 to Kp6 make-up the *K. pneumoniae* complex. Currently, the phylogroups can be reliably identified only by gene sequencing. Misidentification using standard methods is common and the clinical significance of *K. pneumoniae* complex members is therefore imprecisely defined. Here, we evaluated the potential of MALDI-TOF mass spectrometry to discriminate *K. pneumoniae* complex members. We report for the first time the existence of mass spectrometry biomarkers associated with the phylogroups, with a sensitivity and specificity ranging between 80-100% and 97-100%, respectively. Strains within phylogroups Kp1, Kp2, Kp4 and Kp5 each shared two specific peaks not observed in other phylogroups. Kp3 strains shared a peak that was only observed otherwise in Kp5. Finally, Kp6 had a diagnostic peak shared only with Kp1. Kp3 and Kp6 could therefore be identified by exclusion criteria (lacking Kp5 and Kp1-specific peaks, respectively). Further, ranked Pearson correlation clustering of spectra grouped strains according to their phylogroup. These results call for incorporation of spectra of all *K. pneumoniae* complex members into reference MALDI-TOF spectra databases, in which they are currently lacking. This advance may allow for simple and precise identification of *K. pneumoniae* and closely related species, opening the way to a better understanding of their epidemiology, ecology and pathogenesis.

## INTRODUCTION

*Klebsiella pneumoniae* is an increasingly challenging human bacterial pathogen, causing hospital or community-acquired infections that are associated with high rates of antibiotic resistance (1, 2). Population diversity studies have shown that *K. pneumoniae* is in fact part of a complex of species, being phylogenetically closely related to *K. quasipneumoniae* (subsp. *quasipneumoniae* and subsp. *similipneumoniae)* and *K. variicola* (3–5). Before recent taxonomic updates (6, 7), *K. pneumoniae* and the other above taxa were designed as *K. pneumoniae* phylogroups Kp1, Kp2, Kp4 and Kp3, respectively (8). Together with two novel phylogroups (Kp5 and Kp6) described recently (5), these taxa constitute the *K. pneumoniae* complex. Although *K. pneumoniae* is numerically the major cause of human infections among members of the complex, the involvement of the other members of the complex in human infections is gaining recognition (4, 8–12). However, the unsuitability of traditional clinical microbiology methods to distinguish species within the complex leads to high rates of misidentifications (most often as *K. pneumoniae*) that are masking the true clinical significance of each phylogroup and their potential epidemiological specificities (8, 9, 12, 13). In fact, the different members of the *K. pneumoniae* complex can be reliably identified only based on gene sequencing (e.g. *bla*_LEN_, *bla*_OKP_, *bla*_SHV_, *rpoB*, *gyrA*, *parC*) (4, 7, 14). Some PCR-based identification methods were developed but they are prone to errors or do not distinguish all phylogroups (8, 15–17). Clearly, there is a need for reliable, cost-effective and fast identification methods able to discriminate members of the *K. pneumoniae* complex.

Matrix-assisted laser desorption ionization-time of flight (MALDI-TOF) mass spectrometry (MS) has revolutionized routine identification of microorganisms, being a fast and cost-effective technique. It now represents a first line identification method in many clinical, environmental and food microbiology laboratories (18). In the case of the *K. pneumoniae* complex, MALDI-TOF MS identification remains largely unsatisfactory given the absence of well characterized, representative members of the complex in spectral databases. Currently, only *K. pneumoniae* and *K. variicola* are included in the Bruker database (https://www.bruker.com/fileadmin/user_upload/1-Products/Separations_MassSpectrometry/MALDI_Biotyper/US_CA_System/MBT_list_of_organisms10_2017.pdf), and identification of even these two species is imprecise given the lack of reference spectra of other phylogroups (13, 19). To address this important limitation of currently MALDI-TOF MS technology, we used a collection of well characterized reference strains from the six *K. pneumoniae* complex phylogroups and analyzed them by MALDI-TOF MS in order to define the potential of this method to identify species within the *K. pneumoniae* complex.

## MATERIAL AND MEHODS

### Bacterial strains

A set of 46 strains previously characterized by whole-genome sequencing or using core gene sequences (5, 7, 20, 21) were analyzed in this study (Table S1). The strains belonged to the taxa *K. pneumoniae* (*sensu stricto*, i.e., Kp1; n=10), *K. quasipneumoniae* subsp. *quasipneumoniae* (Kp2, n=9), *K. quasipneumoniae* subsp. *similipneumoniae* (Kp4, n=7), *K. variicola* (Kp3, n=9), and to two taxonomically undefined lineages named Kp5 (n=6) and Kp6 (n=5). Strains had been stored in brain heart infusion broth containing 25% glycerol at -80°C and were subcultivated before use in this study.

### Spectra acquisition

An overnight culture on Luria-Bertani agar (37°C, 18h) was used to prepare the samples with the ethanol/formic acid extraction procedure following the manufacturer recommendations (Bruker Daltonics, Bremen, Germany). Samples (1 μL) were spotted onto an MBT Biotarget 96 target plate, air dried and overlaid with 1 μL of a saturated α-cyano-4-hydroxycinnamic acid (HCCA) matrix solution in 50% of acetonitrile and 2.5% of trifluoroacetic acid. Mass spectra were acquired on a Microflex LT mass spectrometer (Bruker Daltonics, Bremen, Germany) using the default parameters (detection in linear positive mode, laser frequency of 60 Hz, ion source voltages of 2.0 and 1.8 kV, lens voltage of 6 kV) within the mass range of 2,000-20,000 Da. For each strain, a total of 24 spectra from 8 independent spots were acquired (3 spectra *per* spot, instrumental replicates). External calibration of the mass spectra was performed using Bruker Bacterial Test Standard (BTS).

### Spectra analysis

The spectra were preprocessed by applying the “smoothing” and “baseline subtraction” procedures available in FlexAnalysis software (Bruker Daltonics, Bremen, Germany), exported as peak lists with *m/z* values and signal intensities for each peak in text format, and imported into a dedicated BioNumerics v7.6 (Applied Maths, Ghent, Belgium) database. Peak detection was performed in BioNumerics using a signal to noise ratio of 20. The instrumental replicates (24 spectra for each strain) were used to generate a mean spectrum for each strain using the following parameters: minimum similarity, 80%; minimum peak detection rate, 60%; constant tolerance, 1; and linear tolerance, 300 ppm. Finally, peak matching was performed to search all distinct peaks (called peak classes in BioNumerics) using as parameters: constant tolerance, 1.9; linear tolerance, 550 ppm; maximum horizontal shift, 1; peak detection rate, 10. The discriminating value of each resulting peak was evaluated by a Mann-Whitney test (22). To allocate proteins associated with peaks, the online tool TagIdent was used (http://web.expasy.org/tagident/). Additionally, a Neighbor Joining tree based on ranked Pearson coefficient was constructed using BioNumerics.

## RESULTS AND DISCUSSION

Forty-six reference strains representing the six phylogroups currently known within the *K. pneumoniae* complex were analyzed by MALDI-TOF MS. Based on the MALDI Biotyper Compass database version 4.1.80 (Bruker Daltonics, Bremen, Germany), the 46 strains were identified either as *K. pneumoniae* (31 strains, all belonging to Kp1, Kp2, Kp4 and Kp6) or as *K. variicola* (15 strains, all strains of Kp3 and Kp5). Identification scores ranged between 2.16-2.56 for *K. pneumoniae* and 1.89-2.55 for *K variicola.* Of note, in two cases a replicate was reported in one measure as *K. pneumoniae* and in other as *K. variicola*. These data highlight the need to update the database in order to refine confidence in *K. pneumoniae*/*K. variicola* identification and to enable identification of *K. quasipneumoniae and novel phylogroups*. **Fig. 1** summarizes the peak positions found in each strain. Most (about 97%) of the peaks were concentrated in the region below 10,000 *m/z* and almost no peak was found above this value. The similarity among spectra within the *K. pneumoniae* complex was always above 87% (data not shown), with peaks at 4363, 5379, 6286, 6298, 7241 and 9473 *m/z* being found in all the members of the complex. Interestingly, ten specific biomarkers associated with specific members of the *K. pneumoniae* complex were identified. These peaks were located within the range 3835 - 9553 *m/z.* The specificity and sensitivity of their distribution among phylogroups ranged between 97-100% and 80-100%, respectively (**Fig. 1 and Table 1**). Kp1 (4152 and 8305 *m/z*), Kp2 (4136 and 8271 *m/z*), Kp4 (7670 and 3835 *m/z*) and Kp5 (4777 and 9553 *m/z*) each presented two specific peaks, which may allow their unambiguous identification. Kp3 strains shared a peak that was only observed otherwise in Kp5 (7768 *m/z*). Finally, Kp6 had a diagnostic peak (5278 *m/z*) shared only with Kp1. Kp3 and Kp6 could therefore be identified by exclusion criteria (lacking Kp5 and Kp1-specific peaks, respectively) (**Fig. 1 and Table 1**). These data reveal the possibility to identify precisely an isolate of the Kp complex based on the specific combination of the above described peaks. To the best of our knowledge, this is the first time that mass spectrometry biomarkers that discriminate the phylogroups of the *K. pneumoniae* complex are described. Furthermore, cluster analysis grouped all strains according to their phylogroup (**Fig. S1**), also showing the potential of whole spectrum comparison for strain identification at the phylogroup level.

**Figure 1.**
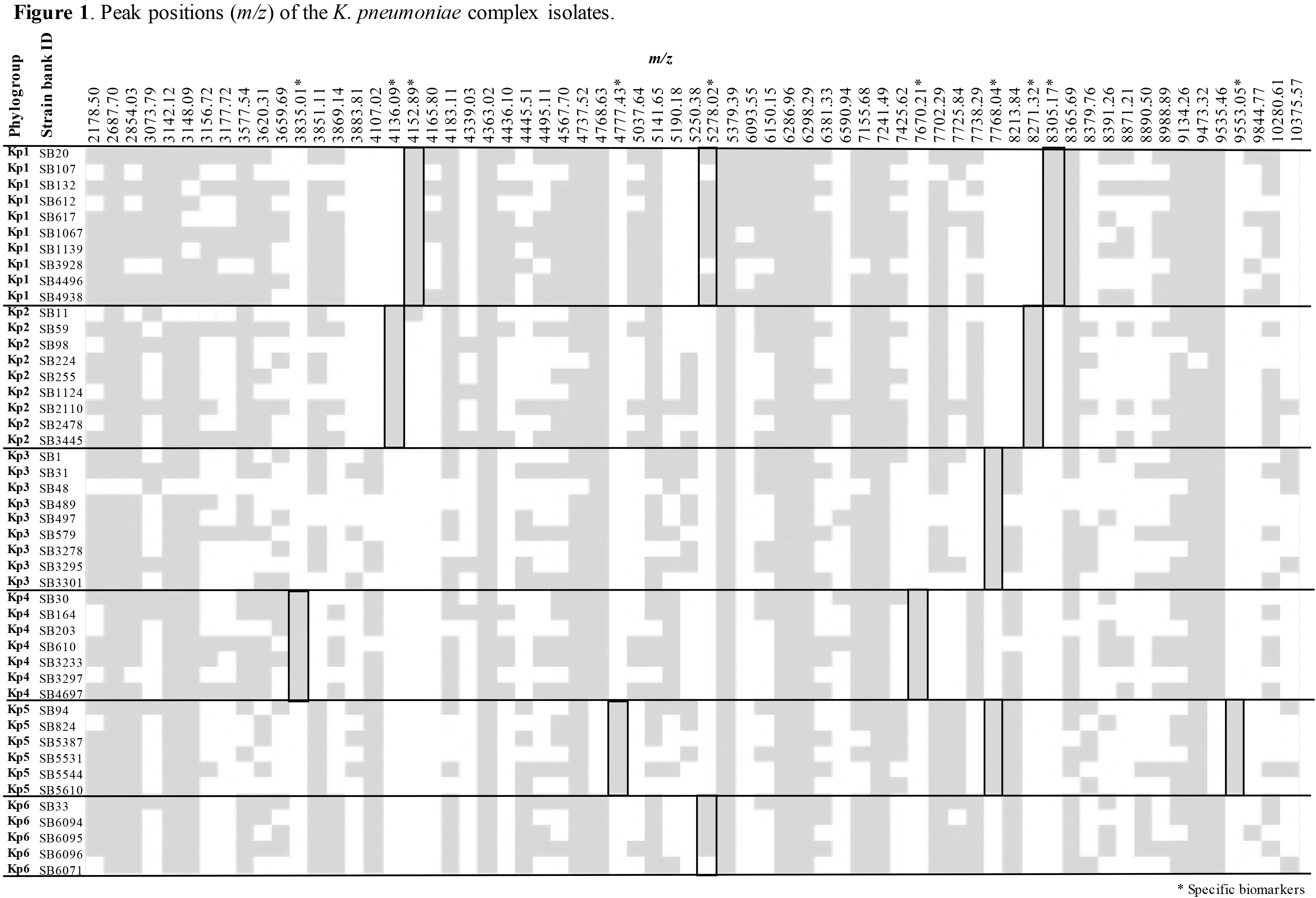
Peak positions (*m/z*) for each of the *K. pneumoniae* complex strains. Star denotes those peaks that are useful for discrimination among phylogroups, as detailed in Table 1.

**Table 1.** MALDI-TOF mass spectrometry peak biomarkers useful to discriminate *Klebsiella pneumoniae* phylogroups

| **Present in Kp phylogroup(s):** | **Peak Position**<br>**(*m/z*)<sup>1</sup>** | **Sensitivity** | **Specificity** | **Possible proteins<sup>2</sup>** |
| --- | --- | --- | --- | --- |
| **Kp1** | 4152.89 | 100% | 97.3% | - |
|  | 8305.17 | 100% | 100% | - |
| **Kp2** | 4136.09 | 100% | 100% | - |
|  | 8271.32 | 100% | 100% | - |
| **Kp3 and Kp5** | 7768.04 | 100% | 100% | Ribosomal protein L31 |
| **Kp4** | 3835.01 | 100% | 100% | - |
|  | 7670.21 | 100% | 100% | Uncharacterized protein<br>specific for Kp4 |
| **Kp5** | 4777.43 | 100% | 100% | - |
|  | 9553.05 | 100% | 100% | - |
| **Kp1 and Kp6** | 5278.02 | 80% | 100% | Ribosomal protein S22 |

<sup>1</sup> Position in the spectra using a tolerance of $\pm 0.05\%$ .
<sup>2</sup> As determined using TagIdent.

About half of the peaks visualized in a bacterial spectrum in the mass range used in this work (2,000-20,000 Da) correspond to ribosomal proteins (18). Here, we were able to presumptively identify two of the specific peaks as ribosomal proteins (S22 and L31, respectively 5278 and 7768 *m/z*), and one as a non-characterized protein specific for Kp4 [7670 *m/z*, locus tag SB30_RS24725 (GenBank Accession number CBZR010000000)]. The specificity of the peaks was supported by the protein alignments obtained from whole-genome sequences (data not shown). The other seven peaks useful for identification could not be associated with a defined protein (**Table 1**).

In conclusion, this work demonstrates the potential of MALDI-TOF MS to identify isolates of the *K. pneumoniae* complex at the phylogroup level. We urge that reference spectra of the various taxa of the *K. pneumoniae* complex be incorporated into reference MALDI-TOF spectra databases, so that the approach could be implemented in microbiology laboratories. Improved identification of *K. pneumoniae* and related taxa will advance our understanding of the epidemiology, ecology and links with pathogenesis of this increasingly important group of pathogens.

## Acknowledgments

We thank the Biological Resource Center of Institut Pasteur (Chantal Bizet and Estelle Muhle) for support within the MALDI-TOF MS platform. We acknowledge Andriniaina Rakotondrasoa and Jean-Marc Collard (Institut Pasteur of Madagascar), S. Wesley Long (Houston Methodist Hospital) and David A. Rasko and J. Kristie Johnson (University of Maryland School of Medicine, Baltimore) for kindly providing strains analyzed in this study.

This work is part of the MedVetKlebs project, a component of the One Health European Joint Programme, which has received funding from the European Union’s Horizon 2020 research and innovation programme under Grant Agreement No 773830.

